# Avian predator-prey dynamics in a changing climate along the Western Antarctic Peninsula; a scoping review

**DOI:** 10.1101/2024.10.14.618217

**Authors:** Tamara M. Russell, Victoria R. Hermanson

## Abstract

A unique characteristic of the food web along the Western Antarctic Peninsula (WAP), one of the fastest warming regions in the world, is that the avian tertiary predators seasonally rely on avian secondary predators for their subsistence. We conducted a scoping review to 1.) provide a summary of research on Antarctic avian predator-prey relationships, 2.) investigate potential avian predator-prey relationships and trends with the environment, and 3.) highlight research gaps and provide recommendations for future research. We searched Web of Science and Google Scholar for publications in English during any years. For our first aim, we searched using the terms “predator-prey dynamics” AND “Antarctica.” We excluded results that did not include both avian predators and prey, which resulted in eight publications from around the Southern Ocean, and one along the WAP. For our second aim, we searched using the terms of each species’ common and scientific names (gentoo penguin, *Pygoscelis papua*, Adelie penguin, *P. adeliae*, chinstrap penguin, *P. antarcticus*, southern giant petrel, *Macronectes giganteus*, south polar skua, *Stercorarius maccormicki*, brown skua*, S. antarcticus*) AND “population” AND “Antarctic Peninsula.” We refined our results (N=59) to publications with data on at least one prey and one predator avian species of all papers found in Web of Science, and the first 100 records of Google Scholar. We selected five locations that had data spanning over 10 years and that spread across the northern WAP. We compared predator-prey species trends across time along with sea surface and air temperature. We found that predator-prey dynamics between avian secondary and tertiary predators have had limited investigations in Antarctica. Along the WAP, the relationship between different penguin species and avian tertiary predators are highly variable and many population trends are decoupled from local temperature change. We include recommendations for future data collection and research on these interactions.

## INTRODUCTION

Anthropogenic-driven warming is affecting marine ecosystems through multiple processes, including direct sea surface and air warming, increased stratification, and decreases in sea ice formation [1, 2]. These environmental changes may affect marine organisms through multiple processes, including through impacts on habitat suitability and prey availability [1, 3], and can result in shifts in the phenology, distribution, and predation intensity of species [2, 4, 5]. Climate change driven decreases and/or range shifts of prey species can negatively affect predator populations, alternatively, decreases in prey can make predation disproportionately impactful on prey populations and reproductive success [3]. Such shifts in predator-prey interactions may have ecosystem-level consequences, therefore understanding these dynamics and their relationships to environmental change is critical for conservation and management of marine systems [6, 7].

Polar ecosystems are already experiencing effects of climate change [2, 8]. In the Southern Ocean, the Western Antarctic Peninsula (WAP) is one of the fastest warming regions in the world and rapidly experiencing effects of climate change, including increasing ocean and air temperatures, changes in wind patterns, and the extent, thickness and seasonality of sea ice, a critical characteristic of this habitat [9, 10, 11]. Along the WAP, the sea ice duration has already shortened by approximately 3 days per year and the sea ice extent has decreased by 5-6% per decade [10]. Sea ice characteristics are critical to this ecosystem, as it provides habitat, food through algal growth, breeding platforms, and can affect the success and survival of many Antarctic species [11, 12, 13]. Because of the magnitude of change and species-specific adaptations to this environment, this region is projected to be a hotspot for changes in species composition and diversity, including local extinctions and invasions [14, 15, 16].

There is an abundance of avian marine predators along the WAP, including secondary predators, such as brush-tailed penguins (*Pygocelis*) that primarily feed on Antarctic krill, and tertiary predators such as south polar (*Stercorarius maccormicki*) and brown skuas (*S. antarcticus*), and southern giant petrels (*Macronectes giganteus*). These tertiary predators feed on a variety of foods, including penguins (Fig 1). There are three species of Pygocelids that breed and feed along the WAP: chinstrap (*Pygoscelis antarcticus*), Adélie (*P. adeliae*), and gentoo (*P. papua*). Pygocelids primarily feed on krill, although gentoo penguins are more generalist and occasionally consume fishes and squid [17, 18, 19]. Both the skuas and giant petrels predate upon penguin resources (eggs, chicks, and even adults by the giant petrels), but also rely heavily upon scavenging and at-sea surface feeding of fishes, cephalopods, and invertebrates [20]. Giant petrels in particular are a fierce predator in this ecosystem, and occupy the highest trophic level among birds in the WAP [21]. Skuas are also known for inducing penguins to regurgitate krill or will scavenge on krill that failed to make it into a chick’s throat (i.e., krill spills; [22]). Brown skuas in particular have a predominantly predatory lifestyle, focusing on penguin resources, placentae, and carrion [23], with limited offshore foraging [24]. While, south polar skuas consume a high proportion of fish [25], and are regularly seen foraging at sea [26, 27].

**Fig 1:**
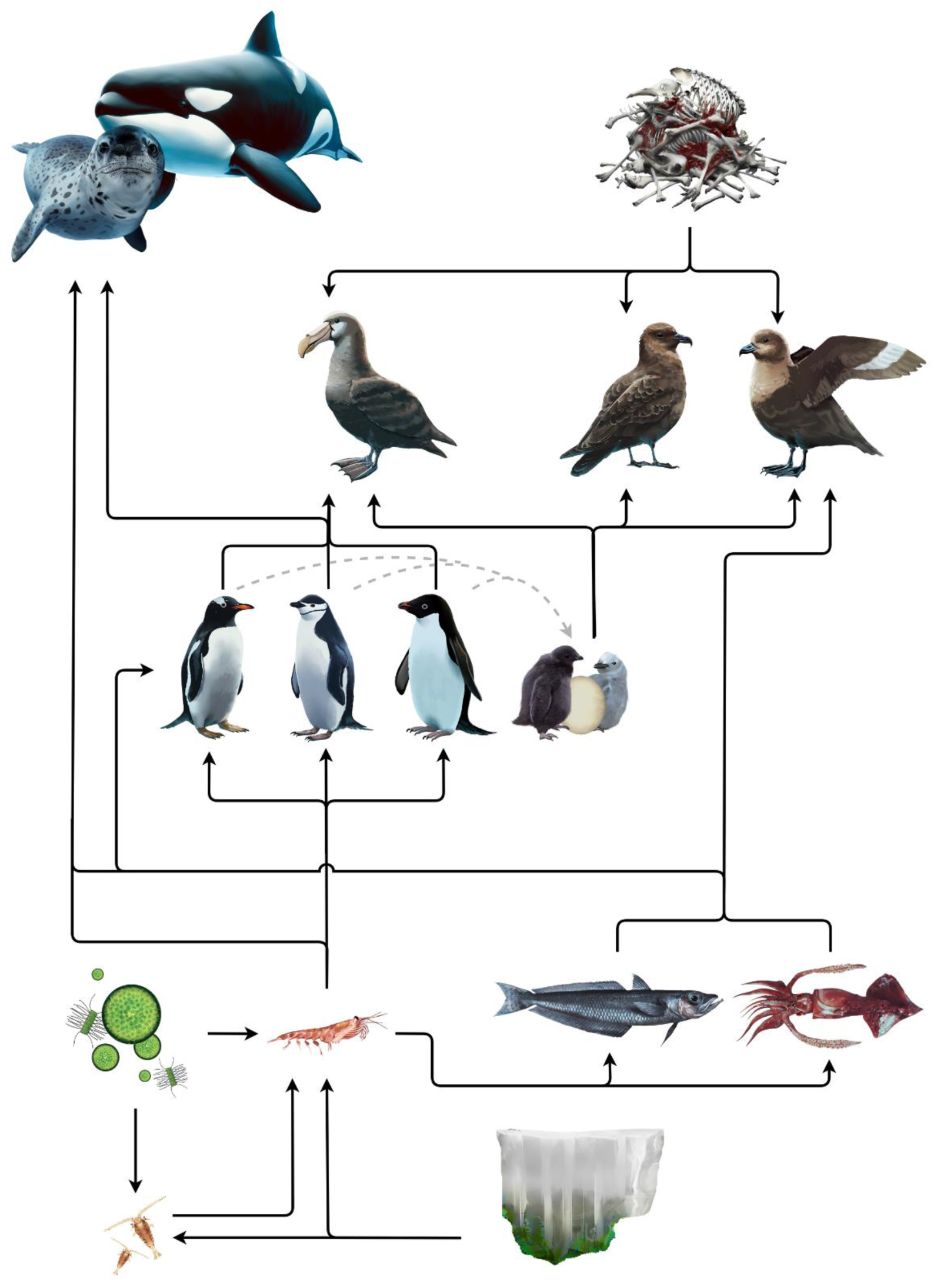
Simplified Western Antarctic Peninsula (WAP) food web, focused on connections with seabird species (© Freya Hammar). Sea ice algae and pelagic phytoplankton are displayed as separate components, although there may be overlap in species. This is to highlight the importance of sea ice in this ecosystem, as increases in sea surface and air temperatures increase there are impacts on sea ice along the WAP.

Much of the long-term monitoring and avian research in the WAP has focused on the brush-tailed penguin species, and changes in their population success and distribution have been documented in association with ocean warming and sea ice conditions [28, 29, 30]. In this region, Adélie penguins are declining, chinstrap penguins show variable responses but overall declines, and gentoo penguins, a generalist that needs ice free conditions, have been increasing [28, 31, 32, 33]. However, the population trends of avian tertiary predators remains under- studied, and these energy pathways are typically left out of food web models (e.g., [34, 35]). It is unknown whether shifts in the environment can also influence penguin species indirectly, through changes in interspecies interactions with predatory seabirds. With colonies of large penguin populations, these interactions may not be readily apparent or may have an insignificant impact on penguin numbers or reproductive success. As species decline, as is the case for many chinstrap and Adélie populations along the WAP, the role of predators may play a larger role in their success and population dynamics.

Because ecosystem-based management is the primary driver in Antarctic resource management [36, 37], it is important to consider ecosystem interactions that are not currently represented. While predator-prey interactions between upper trophic level predators and lower trophic level prey [38, 39, 40] are at the forefront of ecosystem-based management research, interactions between upper trophic level predator-prey can provide critical insight into how the Antarctic food web is evolving with a changing climate. In this paper, we use a qualitative to investigate WAP Antarctic avian predator dynamics by addressing these three goals: 1) review published literature on Antarctic avian predator-prey dynamics; 2) investigate these species population trends and relationships with environmental conditions; and 3) provide recommendation for future research priorities.

This work is a first-step in better understanding the complexities of avian predator interactions in this region, and we hope to inspire future work in order to better monitor and predict future changes within this vulnerable ecosystem.

## METHODS

### Protocol and search strategy

To conduct these reviews, we used Web of Science, and then a supplemental search with Google Scholar using the same terms. Both authors worked independently on different search terms and tracked results in a shared Google Sheet, and followed the same protocol to reduce bias. Our scoping review protocol included searching for articles in English across all years and the combination of search terms could appear anywhere in the article; searches began on 07 July 2023 and ended on 23 September 2023. To address our first research aim of this study, to review current knowledge of Antarctic avian predator-prey dynamics, we searched for predator-prey information, including the terms “predator-prey dynamics” and “Antarctic Peninsula.” To address our second research aim, to investigate species trends and relationships with the environment, we searched for avian predator and prey time-series data that extended across at least 10 years, during any time-period, collected from colonies along the WAP. The search terms included both the species’ common and scientific names of the six species (Fig 1), “population” AND “Antarctic Peninsula.”

### Inclusion and exclusion criteria

After conducting our literature searches, we refined results to specifically address our two aims; providing an overview of research on Antarctic predator-prey dynamics and identify time series for comparisons between predators and prey along the WAP. We refined our results by manually evaluating the title, keywords and/or abstracts of all papers found in Web of Science, and the first 100 records of Google Scholar searches. For our first aim, author T.M.R. selected publications if they compared an avian predator and avian prey, and if it was research from Antarctica. For our second aim, we filtered our results by evaluating article titles and abstracts to decide whether it contained Antarctic seabird time series. From these results, we then extracted data including year, species, and data types. As many of our selected papers utilized the Oceanites Antarctic Site Inventory (OASI), we downloaded data from the OASI (https://www.penguinmap.com/mapppd, [41,42]). Locations were additionally filtered by evaluating whether the data available spanned more than 10 years, and if there were data on both avian prey (i.e., chinstrap, gentoo, or Adélie penguins) and avian predators (i.e., southern giant petrels, brown and south polar skuas). We then selected our final locations for analysis based on their latitudinal spread and extracted the data for each year, for each species and data type.

### Data synthesis and characteristics

#### Seabird predator-prey data

The selected data used to investigate trends all came from locations within the South Shetland Islands, which are located along the northwestern Antarctic Peninsula (within 60-63°S, 57-62°W; Fig 2). The South Shetland Islands are situated between the Bransfield Strait to the south, the Bellingshausen Sea to the southwest, and the Drake Passage and Southern Ocean in the north.

**Fig 2:**
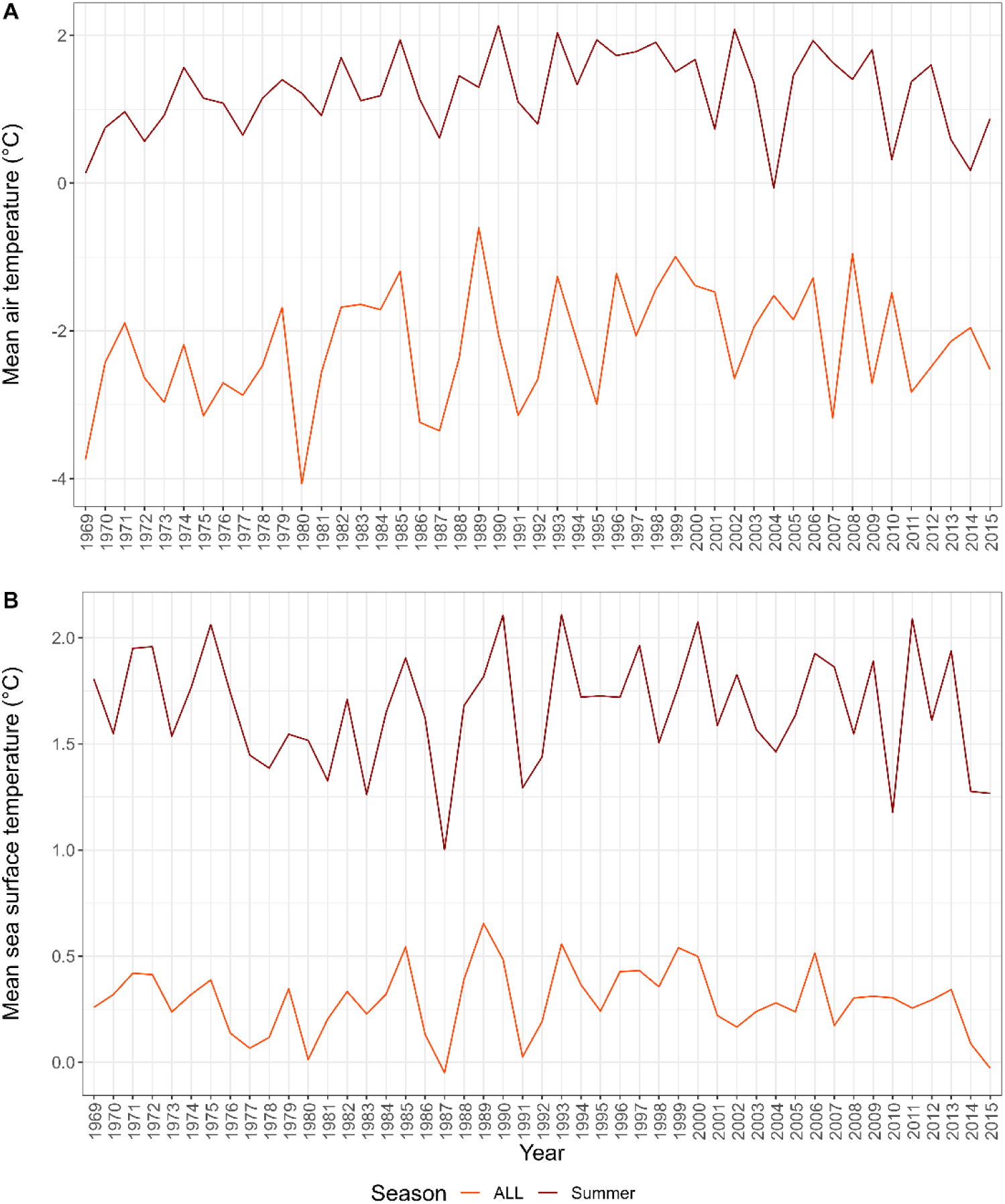
Annual and summer averaged environmental data for the South Shetland Islands and surrounding Southern Ocean. Air temperature data (A) were derived from monthly measurements from five stations around the South Shetland Islands [43]. Sea surface temperature (SST) data (B) were obtained from daily measurements (HadISST 1° daily; [44]). The annual air temperatures have increased in this region (Adj. R^2^= 0.14, F_(1,56)_=9.02, p value=0.004), while there was no significant trend in SST (Adj. R^2^= -0.02, F_(1,47)_=0.007, p-value=0.9).

Data types varied among locations and included the number of breeding pairs, number of active nests, and number of chicks. If more than one datum was collected per year, we used the average in analysis. As Antarctic data are typically collected in the austral summer, it spans across the new year (e.g., Dec 1995 to Jan 1996). We used the later year in our analysis, for example, data collected during the 1995/1996 season is 1996. We used only species-level data in our analysis and discarded grouped data; which was only available for skuas. A grouped ‘SKUA’ term in our selected datasets included records of birds that either could not be identified or were not attempted at being identified to species. As brown and south polar skua have different diets, we wanted to test for potential differences, therefore only using species-level data.

#### Environmental data

We used two environmental variables to test for relationships between species abundance and local and regional conditions between 1968 and 2015. As a proxy for local conditions during the breeding season, we used air temperature data obtained from the READER project ([43]; https://legacy.bas.ac.uk/met/READER). We selected five stations around the South Shetland Islands (Captain Arturo Prat Base, Comandante Ferraz Antarctic Station, Deception Station, Great Wall Station, Carlini Base,King Sejong Station, and Teniente R. Marsh Airport), and took the average summer (December-February) temperature among all stations (Fig 2). For regional, year-round conditions, we used sea surface temperature (SST) data from the Hadley Centre Global Sea Ice and Sea Surface Temperature (HadISST, [44]) obtained from National Center for Atmospheric Research (https://rda.ucar.edu/datasets/ds277.3). The HadISST data is 1° daily means that are constructed by combining observations from different sources (ships, satellites, and buoys), along with reconstruction of historical conditions [44]. SST data were extracted within the bounding box that captured the South Shetland Islands and potential foraging habitat for seabirds that nest there (60° to 64° S, 55° to 66° W). Some of the species (e.g., south polar skua) forage much further outside of this region, however these area captures breeding foraging habitat as well as a portion of wintering foraging habitat for most other species in our study. We calculated the annual average within these gridded data as an estimate of the broad conditions experienced by these species (Fig 2).

### Data analyses

In addition to reviewing previous research on avian predators and prey in the WAP, we compared temporal trends among species at selected colonies to investigate patterns in predator and prey populations, along with relationships with sea surface and air temperatures. These are a first approach in understanding how avian predator and prey population trends may be linked.

For example, we might expect that at large penguin colonies, their fluctuations or trends in abundance may not effect predator populations, whereas at small penguin colonies, fluctuations and/or declines could have an impact on how well and/or abundant predator populations do.

All statistical analysis and plots were produced using R programming [45]. Multiple linear regression models were conducted on these data to test for linear relationships of species annual trends and environmental variable relationships. Data was truncated within the years 1968 to 2016, a time period where both SST and air temperature were available and to only species and data types that had over 4 observations so they could be analyzed in the multiple regression models. To prevent trends that could have resulted from interannual variability, all time series used in these analyses spanned at least 10 years, although we recognize longer time series are needed to fully capture long-term impacts from climate change.

To further compare avian predator and prey trends, we used Pearson’s correlation analysis to directly compare years where both predator and prey had data available and provide additional information on the strength and direction of relationships between predator-prey species. The Pearson correlation coefficient (r) measures how closely related two variables are, from -1 as a perfect negative linear relationship to +1 as a perfect positive linear relationship, and 0 indicates that there is no linear correlation between them. We classified r between 0.40 and 0.60 as moderate and over 0.60 as strong correlations [46]. We calculated and plotted Pearson’s correlation coefficients using the ‘ggpairs’ function in the GGally library in R [47].

## RESULTS

Our literature search on publications on Antarctic avian predator-prey relationship initially yielded 31 results on Web of Science and 206 results on Google Scholar. After filtering our results, we identified eight publications that addressed the relationships of avian predators and prey around the Southern Ocean [48, 49, 50, 51, 52, 53, 54], and one from the WAP [22].

The initial results of our literature search on avian predator and prey time series data were 1,437 results on Web of Science and 6,070 results on Google Scholar. After evaluating all of the Web of Science results and the first 100 for each species on Google Scholar, we had identified 59 articles with time series data on our species of interest. After further evaluation using our selection protocol we selected 18 publications (Table 1, [55:71]) that included data from five locations from around the South Shetland Islands for analysis based on their latitudinal spread and extracted the data for each year, for each species and data type (Fig 3).

**Fig 3:**
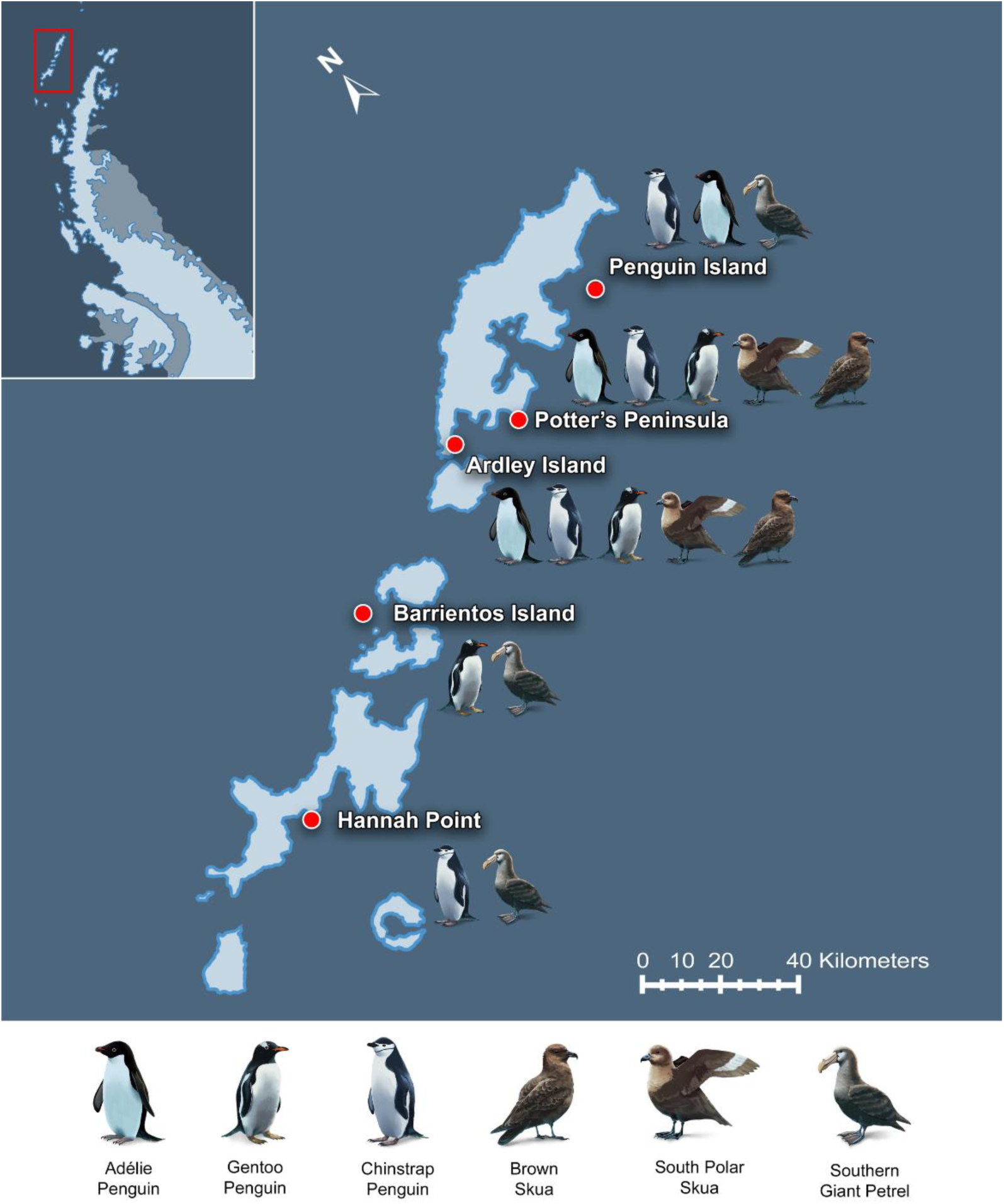
Map of the locations of the data that were extracted from published reports and analyzed in this study. Alongside each location, are the species that had data available and were used in this study. Due to lack of multi-species, publicly available data, our review did not extend beyond the South Shetland Islands (© Freya Hammar).

**Table 1:** Description of the data used in this study, including the location of the colony, the species and type of data, the range of years data was available and the sources we acquired the data from.

| Site | Species | Data type | Year Range | Source |
| --- | --- | --- | --- | --- |
| Penguin Island | CHPE | Breeding pairs | 1979-2013 | [55, 56, 57, 58, 59, 60, |
| Penguin Island | ADPE | Breeding pairs | 1979-2013 | [55, 56, 57, 58, 59, 60, |
| Penguin Island | SGPE | Breeding pairs | 1979-2012 | [55, 57, 58, 61] |
| Potter Peninsula | BRSK, SPSK | Breeding pairs | 1989-2014 | [62, 63, 64, 65, 66] |
| Potter Peninsula | ADPE | Breeding pairs | 1972-2013 | [67, 68] |
| Potter Peninsula | GEPE | Breeding pairs | 1972-1989 | [67] |
| Potter Peninsula | CHPE | Breeding pairs | 1972-1989 | [67] |
| Potter Peninsula | ADPE | Nests | 1996-2011 | [67, 68, 69] |
| Potter Peninsula | GEPE | Nests | 1997-2007 | [67] |
| Potter Peninsula | ADPE | Chicks | 1996-2007 | [70] |
| Potter Peninsula | GEPE | Chicks | 1997-2007 | [70] |
| Ardley Island | GEPE | Nests | 1980-2005 | [71] |
| Ardley Island | CHPE | Nests | 1980-2005 | [71] |
| Ardley Island | ADPE | Nests | 1980-2012 | [71] |
| Ardley Island | BRSK, SPSK | Breeding pairs | 1980-2015 | [62, 65, 69, 72] |
| Barrientos Island | SGPE | Nests | 2000-2012 | [71] |
| Barrientos Island | GEPE | Nests | 2000-2012 | [71] |
| Hannah Point | CHPE | Nests | 1987-2011 | [71] |
| Hannah Point | SGPE | Nests | 1998-2011 | [71] |

\*Species: Chinstrap (CHPE), Adelie (ADPE), and gentoo (GEPE) penguins, South polar (SPSK), brown (BRSK), and southern giant petrels (SGPE).

### Trends among avian predators and prey species

Time-series results within sites varied (Table 2, Fig 4). In the north, breeding pairs of chinstrap and Adélie penguins on Penguin Island have both declined, but our model could not significantly predict these trends; while giant petrel breeding pairs on Penguin Island show no trend. Further south, at Potter’s Peninsula, there were declines in Adélie breeding pairs, nests, and chicks (R^2^=0.90, p<0.0001, R^2^=0.86, p<0.0001, and R^2^=0.75, p=0.003, respectively), although environmental variables were not significant predictors of these trends. There were no apparent trends in chinstrap or gentoo penguins or either species of skua, or any correlations with environmental variables at Potter’s Peninsula. Continuing southeast to Ardley Island, both chinstrap and Adélie nests are declining and with year as the only significant variable for these two species (R^2^=0.61, p=0.0006 and R^2^=0.59, p<0.002, respectively). In contrast, gentoo penguin nests here are increasing, but not driven by environmental conditions (R^2^=0.45, p=0.010). Our models did not explain the trends for brown or south polar skuas at Ardley Island, although the year variable was significant (p=0.024). On Barrientos Island in the south, giant petrel nests have significantly declined, with the variability explained by year (R^2^=0.32, p=0.048) and no environmental conditions. Although there is an apparent increase in gentoo penguins on Barrientos (Fig 4), these trends are not significant. At Hannah Point, chinstrap penguin nests have decreased here with both year explaining a significant about of this trend (R^2^=0.75, p=0.037). The full model for southern giant petrels at this colony was not significant, however, year was a significant variable (p=0.016).

- **Table 1**: Multiple linear regression results between species at each colony with Year, sea surface (SST) and air temperature; including
- the number of years (n), model results, and results for independent variables. Significant trends (p<0.05) are in bold.
- *Species: Chinstrap (CHPE), Adelie (ADPE), and gentoo (GEPE) penguins, South polar skua (SPSK), brown skua (BRSK), and
- southern giant petrels (SGPE).

**Fig 4:**
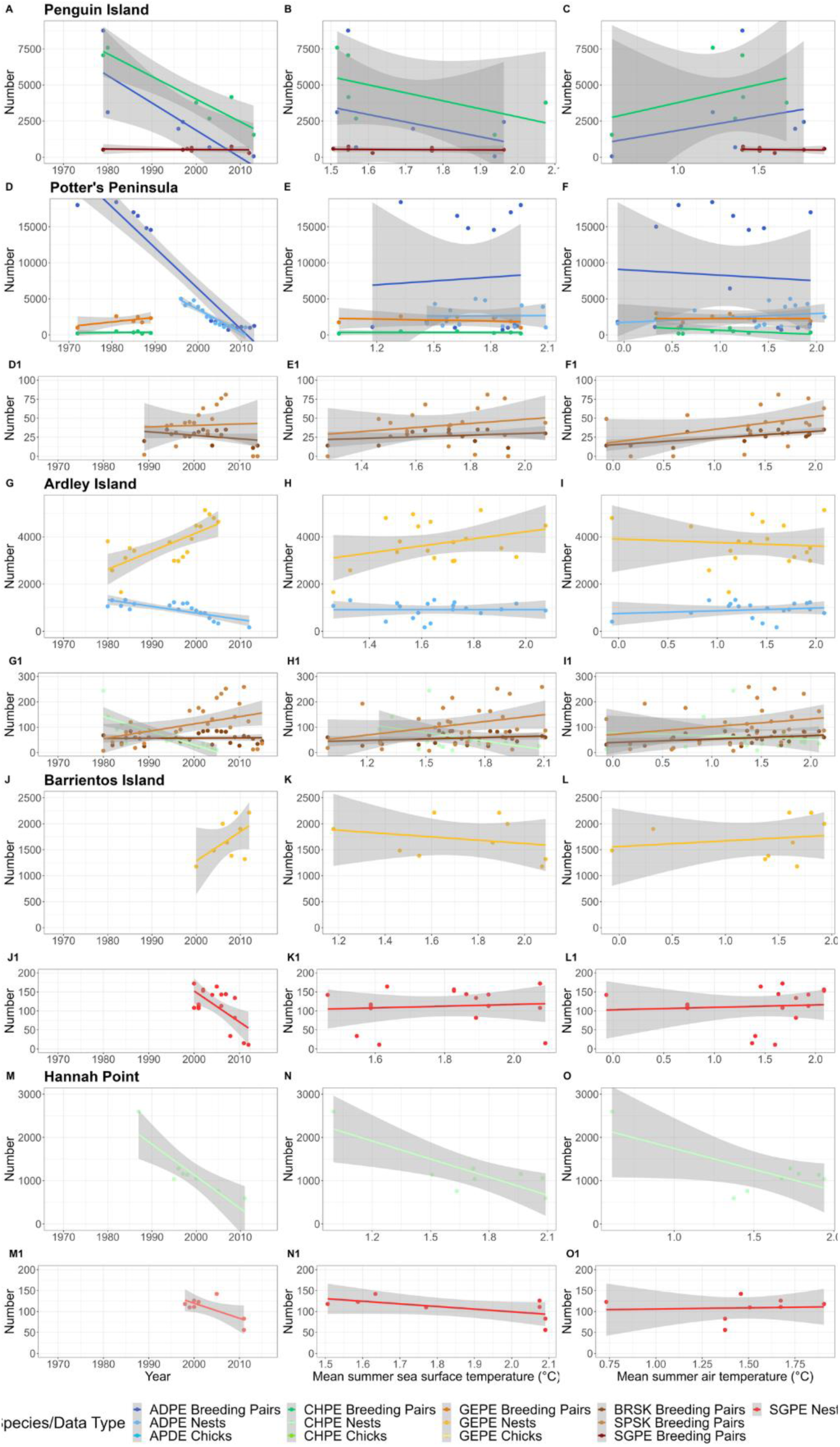
Time series plots of seabird data used in this study (Table 1). The data types varied (number of breeding pairs, chicks, and nests). In order to view these together, we took the log of all counts. Species includes Chinstrap (CHPE), Adelie (ADPE), and gentoo (GEPE) penguins, South polar skua (SPSK), brown skua (BRSK), and southern giant petrels (SGPE).

**Table 2:**
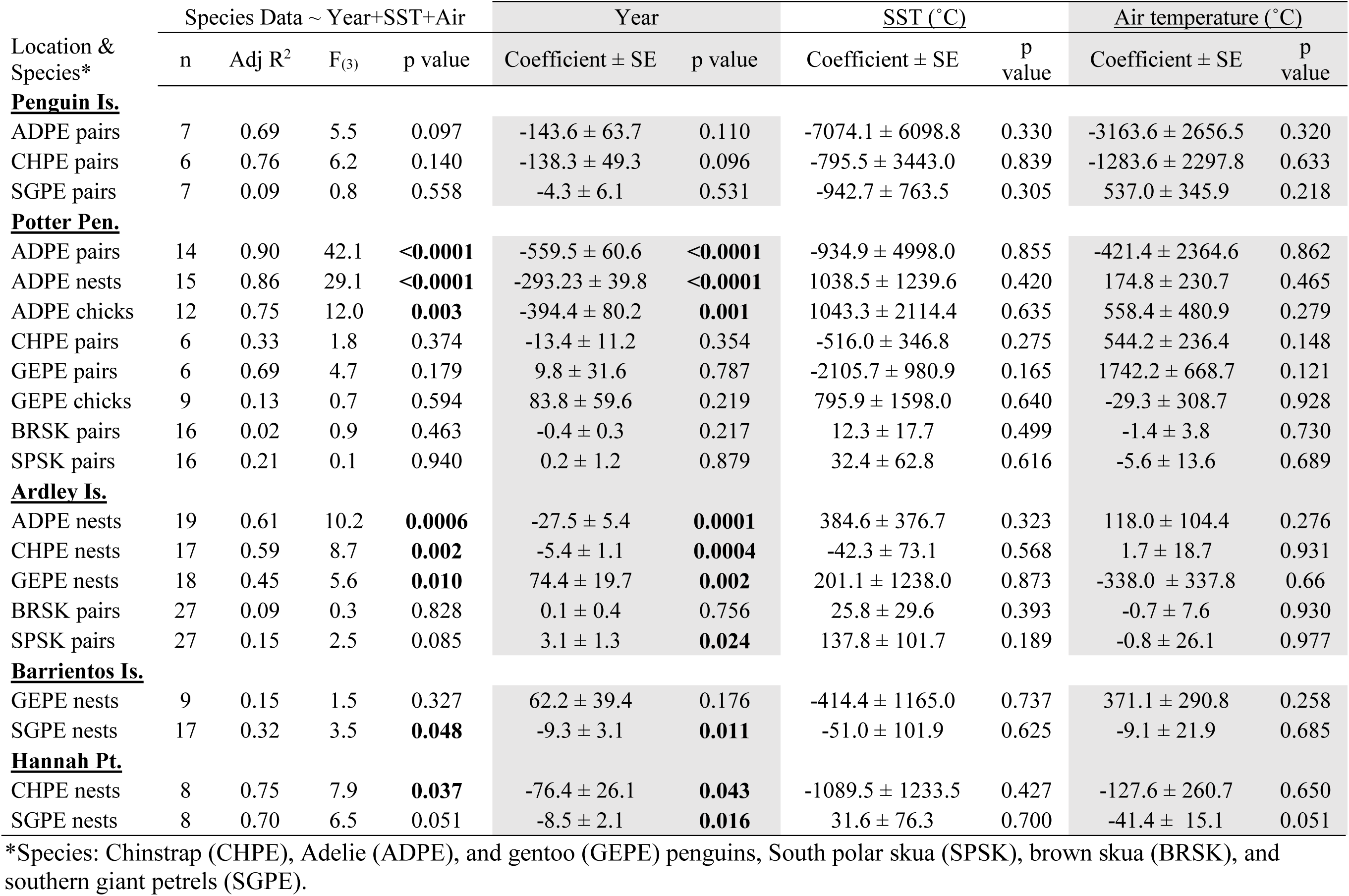
Multiple linear regression results between species at each colony with Year, sea surface (SST) and air temperature; including the number of years (n), model results, and results for independent variables. Significant trends (p<0.05) are in bold.

Pearson’s correlation coefficients varied between species and data types among sites (Table 3, Fig 1S-8S). On Penguin Island, there was a strong, negative relationship between Adélie and giant petrel breeding pairs (-0.662). At Potter’s Peninsula, Adélie breeding pairs had a moderate negative relationship (-0.555) and Adélie chicks had a strong, negative relationship (- 0.778) with south polar skua breeding pairs not neither with brown skuas. On Ardley Island, both Adélie and chinstrap penguin nests had a moderate to strong, negative relationship (-0.565 and - 0.782, respectively) with south polar skua breeding pairs. However, gentoo penguin nests had a strong, positive relationship (0.609) with south polar skuas and a moderate, positive relationship (0.446) with brown skua breeding pairs. There was no relationship between gentoo penguin and southern giant petrel nests at Barrientos Island. At Hannah Point, chinstrap penguin nests had a strong, positive relationship with giant petrel nests (0.621).

**Table 3:** The relationships, measured by Pearson’s correlation coefficient (r) between avian prey species (CHPE, ADPE, GEPE*) and avian predator species (BRSK, SPSK, SGPE*) in this study. This analysis was only done between species and data types that had three or more data points from the same year. We classified r between 0.40 and 0.60 as moderate (italicized) and over 0.60 as strong (bold) correlations. The Pearson correlation coefficient measures how closely related two variables are, from -1 as a perfect negative linear relationship to +1 as a perfect positive linear relationship, and 0 indicates that there is no linear correlation between them.

| Site | Prey species | Data type | Predator species | Data type | Pearson's $r$ |
| --- | --- | --- | --- | --- | --- |
| Penguin Island | ADPE | Breeding pairs | SGPE | Breeding pairs | **-0.662** |
| Potter's Peninsula | ADPE | Breeding pairs | BRSK | Breeding pairs | -0.174 |
| Potter's Peninsula | ADPE | Breeding pairs | SPSK | Breeding pairs | -0.555 |
| Potter's Peninsula | ADPE | Nests | BRSK | Breeding pairs | -0.087 |
| Potter's Peninsula | ADPE | Nests | SPSK | Breeding pairs | **-0.778** |
| Ardley Island | GEPE | Nests | BRSK | Breeding pairs | *0.446* |
| Ardley Island | GEPE | Nests | SPSK | Breeding pairs | **0.609** |
| Ardley Island | CHPE | Nests | BRSK | Breeding pairs | -0.067 |
| Ardley Island | CHPE | Nests | SPSK | Breeding pairs | **-0.782** |
| Ardley Island | ADPE | Nests | BRSK | Breeding pairs | -0.236 |
| Ardley Island | ADPE | Nests | SPSK | Breeding pairs | -0.565 |
| Barrientos Island | GEPE | Nests | SGPE | Nests | -0.087 |
| Hannah Point | CHPE | Nests | SGPE | Nests | **0.621** |

\*Species: Chinstrap (CHPE), Adélie (ADPE), and gentoo (GEPE) penguins, South polar skua (SPSK), brown skua (BRSK), and southern giant petrels (SGPE).

## DISCUSSION

Potential dynamics of avian predator and prey species in Antarctica are an understudied area of research in Antarctica. We found limited reports investigating these relationships, and few datasets that had both predator and prey data over common time periods. In addition to these limitations, further data on phenology across species and detailed diet data overtime remains sparse and prohibits detailed insights into the role of avian predators in this unique ecosystem.

### Avian predator-prey dynamics in Antarctica

Previous research on predator-prey dynamics in Antarctica has focused on relationships between Antarctic krill and *Pygocelis* penguins, both abundant components of this unique food web (e.g., [32, 73,74]. Although the diets of avian tertiary predators (e.g., Southern giant petrels and skuas) in this region are relatively well known, there has been limited investigations into the predator-prey dynamics between them and their *Pygocelis* prey. During the breeding season, penguin eggs, chicks, and sometimes adults can make up the majority of these tertiary avian predator’s diets [22, 23, 42]. Previous reports document that most of the known skua and giant petrel colonies are near or within *Pygocelis* colonies, providing easy access to this reliable food source during the breeding season [53, 54, 75, 76]. We found limited investigations into avian predator-prey relationships within the Antarctic region, with most of the research focusing on skua-Adélie relationships around East Antarctica. In this region, there is a significant relationship between south polar skuas and Adélie penguin populations; locations where both are decreasing [48] or both are increasing [52]. At larger Adélie colonies, there are a larger number, but smaller proportion, of chick depredation by south polar skuas and the colonies that had larger numbers of skua territories had the lowest reproductive success [50, 77]. Other research investigated multispecies occurrence including our species of interest, but with the focus of understanding niche overlap and mechanisms that prohibit competitive exclusion among these species [78].

Direct comparisons of multiple species on Elephant Island, located north of the WAP, found that all species of penguin and brown skuas were all declining, while southern giant petrel numbers are increasing [79]. The one study we found that directly compared these avian predator-prey species from the WAP was from King George Island, within the South Shetland Islands [22]. This study documented predation events by skuas and giant petrels on different-sized gentoo and Adélie penguin colonies. They found that small colonies were disproportionately affected by predation and that their reproductive success may be influenced by the size of the colony and the abundance and diversity of avian predators [22], which was similar to other findings from studies in the subantarctic islands [49, 77].

### Relationships between seabird predator-prey trends

Within our five colonies used in this exploratory analyses, many of the penguins displayed trends over time in their abundance, while tertiary predators were highly variable over time. In line with previous published results, at four locations either or both chinstrap and Adélie penguins have decreased, while gentoos are either increasing or stable (Fig 4). The results also varied across regions, for example, Southern giant petrels are stable in the north at Penguin Island, where both penguin species seem to be declining. Penguin Island is a heavily visited area by tourists, and impacts to avian species have been previously noted, including changing nest sites, which may mask actual population trends [59]. However, at both locations in the south (Hannah Point and Barrientos Island) giant petrel nests are declining; with the declines at Hannah Point occurring alongside decreases in the number of chinstrap nests. These differences among sites indicate complex drivers of reproductive success, and potential differences in diets among predators at different sites. With the skua species, we found a variety of trends, with south polar skua increasing overall and brown skua trends in breeding pairs remaining stable.

However, the skua data had high interannual variability, which has been previously report at Potter’s Peninsula colony in both breeding pairs and reproductive success [62].

Where we were able to directly compare predator-prey data per year, there were clear patterns among the relationships between secondary and tertiary avian predators (Table 3). Both ice-reliant species of *Pygocelis* (Adélie and chinstrap) had moderate to strong inverse relationships with both south polar skuas. Whereas, the ice-avoiding, gentoo penguin had a positive relationship with the south polar skuas, and moderate, positive relationship with brown skuas. Although there may be correlations among these trends, south polar skuas are known to eat carrion and ectothermic, pelagic prey, rather than rely on taking live penguins as food as brown skuas do regularly. Therefore, the inverse trends among penguin species with skuas we detected may be due to similarities in the conditions that are good for both gentoo and south polar skuas, which have a negative effect on chinstrap and Adélie penguins. Relationships with southern giant petrels varied, with strong, inverse relationships with Adélie penguins in the north, where the petrel population is stable, and strong, positive relationship with chinstraps in the south, where the population is in decline. On Potter’s Peninsula, the Adélie penguin data provided us the opportunity to compare data types. We found that Adélie breeding pairs and nests displayed the same trends in their relationships with brown and south polar skuas (Table 3), however, the nest data resulted in a stronger negative relationship with south polar skuas. To understand these complex relationships, collecting multiple data types may provide more clarity and should be a priority for continued and future monitoring.

The one publication on avian predator-prey relationships along the WAP highlighted the relationships between prey colony size and predators, with smaller colonies being effected more by predation. When we compare different sized colonies, we found variable patterns between small and large colonies with predator correlations. The chinstrap penguins results are in line with the previous report with the large colony on Hannah Point increasing alongside increases in southern giant petrels, while there was a decrease in the much smaller colony of chinstraps on Ardley Island, which was negatively correlated with south polar skuas. The two colonies with gentoo penguin data for comparisons had similar population numbers, and there were comparable correlations among small to large sized colonies of ADPE with skuas.

As the data are limited, these correlations may be due to a variety of processes, such as similar environmental conditions that benefit generalists, or shifts in foraging locations that affect all species. Without diet data or direct observations of predator amount, we cannot verify proposed relationships. However, the variability in the relationships between penguins and skuas are likely more complex due to their heavy reliance on scavenging, compared to the frequency of direct predation that giant petrels exhibit.

### Relationships between predator-prey dynamics and the environment

In line with previous studies, we found air temperatures had increased around the South Shetland Islands [80], however the SST of the broader region had remained stable within our study period. Although there were significant increases in air temperatures, these data did not explain the species trends. The lack of explanatory power of these environmental variables may be due to the spatial scale we used and/or climatic variability masking overall trends (Fig 2), but may be due to other drivers, such as shifts in prey abundance and availability, which are either not tied directly to temperatures or may have lagged responses.

With the temperature increases that are occurring and/or are projected for this region there will be further impacts to these seabird populations and therefore shifts in the predator-prey relationships between them [81, 82]. For example, Adélie and chinstrap penguins are decreasing in many of these colonies, and if gentoos are not establishing or growing their colonies at those colonies, the predation pressure from skuas and giant petrels may increase disproportionately [22]. Because these tertiary predators are generalists and rely on a wide variety of foods, including scavenging, they may be less affected from the immediate pressures of climate change, at least for now. However, if there are decreases in oceanic prey (i.e., fishes, krill, and squid) it may affect the predation pressure on penguins.

### Data limitations

The lack of fine-scale temporal data prevented more complex analyses and investigations into lagged relationships between predators and prey. More detailed data are necessary for conducting more robust analyses on these relationships. We also recognize that species population and reproductive success are complex, and not purely driven by predator controls.

Our analyses are only correlations and not causes, but form a base for future investigations into these predator-prey dynamics.

We found limited previous research on predator-prey linkages along the WAP, and that tertiary predators (skuas and giant petrels) are often left out of food webs, or that energy pathways from penguins to avian tertiary predators are not incorporated. If flying seabirds are included, they are lumped into a general bird group ignoring the high diversity in foraging strategies and diets which make them incompatible for grouping. The lack of species or functional group resolution in these models results in an incomplete food web assessment.

Penguin population data are used as a metric of prey availability, resource management, and ecosystem monitoring in the WAP, however, potential top-down controls on these penguins are not being incorporated. As certain colonies and/or species decline, these relationships may become more impactful and therefore models missing this component are not capturing the full picture of food web connectivity.

### Recommendations

After this evaluation of predator-prey interactions along with WAP and investigations between these species trends, we have developed a series of recommendations for future research. Firstly, the continuation and expansion of current monitoring programs. The WAP is one of the fastest warming regions on Earth, and as vulnerable secondary consumer species decrease, the proportional effect of predation may increase [22]. Therefore, collecting useful data to monitor the pressure of avian predators will become increasingly important. In addition, as the phenology of these populations change, mismatches between predator-prey interactions may occur [83]. These programs should not only collect data on penguin breeding pairs, but attempts should also be made to collect data on the number of nests and chicks so that a metric of reproductive success and phenology can be monitored. In addition to penguin data, data needs to be collected on any breeding predator species within or around the colonies. The limitations of these data collection are in the ability for researchers to remain at or revisit colonies throughout the breeding season. To confirm these relationships, studies on skua and giant petrel diets overtime within these monitored colonies should be conducted and compared with population data to provide information on how their diets change over time, species preferences of prey, and potential population level impacts on these species [84].

In addition to enhanced and expanded data collection, the use of predator-prey models such as Lotka-Volterra models should be used to further understand these relationships under changing environmental conditions. These tertiary predators and these predator-prey interactions should also be included and with more specificity within food web models to more thoroughly understand the food web dynamics in this region. For example, these species and energy pathways could be included into models such as Ecopath with Ecosim [85] or building specific predator-prey models (e.g., [3]).

## Conclusions

This work highlights the gaps in our ecological understanding of predator-prey dynamics in the WAP food web and how these relationships may shift with climate change. We evaluate potential relationships between secondary and tertiary avian predators and their environment, with variable results. The WAP is one of the fastest warming regions on Earth, and these changes continue to break record temperatures and low sea ice extent [81, 82]. Understanding more of the complexity of these food web interactions will allow us to better manage and project future ecosystem changes. We conclude with recommendations for future research, including data types, to assist with enhanced ecosystem monitoring of this vulnerable ecosystem. Excluding potential effects from top-down controls with monitoring efforts likely results in an incomplete understanding of population dynamics. As these predators are not entirely reliant on penguins as a resource, and are opportunistic, generalists, they may also reflect other food web changes. Just as generalist penguins (e.g., gentoo) are the ecological “winners” with warming temperatures, generalist predators of the Antarctic affected differently by fluctuations in specific species, but be impacted by overall decreases in some colonies [86]. Monitoring both secondary and tertiary consumer data provides a deeper understanding of ecosystem structure and functioning, and important data for use in marine conservation.

## Acknowledgments

We would like to thank all those involved in the data collection on marine birds that has occurred throughout the WAP, including tourists that serve as citizen scientists to collect meaningful data. We also thank B. Jack Pan for his assistance with the environmental data used in this research, and thank Stephanie A. Matthews and Allison M. Cusick for their feedback on this work.

## Availability of data

All data is publicly available and can be found through our listed sources (Table 1).

## Supplementary Materials

Pearson’s correlation coefficient plots between species at each site, calculated and plotted using the ggpairs function in the GGally library in R [1]. The Pearson correlation coefficient (r) measures how closely related two variables are, from -1 as a perfect negative linear relationship to +1 as a perfect positive linear relationship, and 0 indicates that there is no linear correlation between them. We classified r between 0.40 and 0.60 as moderate and over 0.60 as strong correlations [2].

**Figure 1S:**
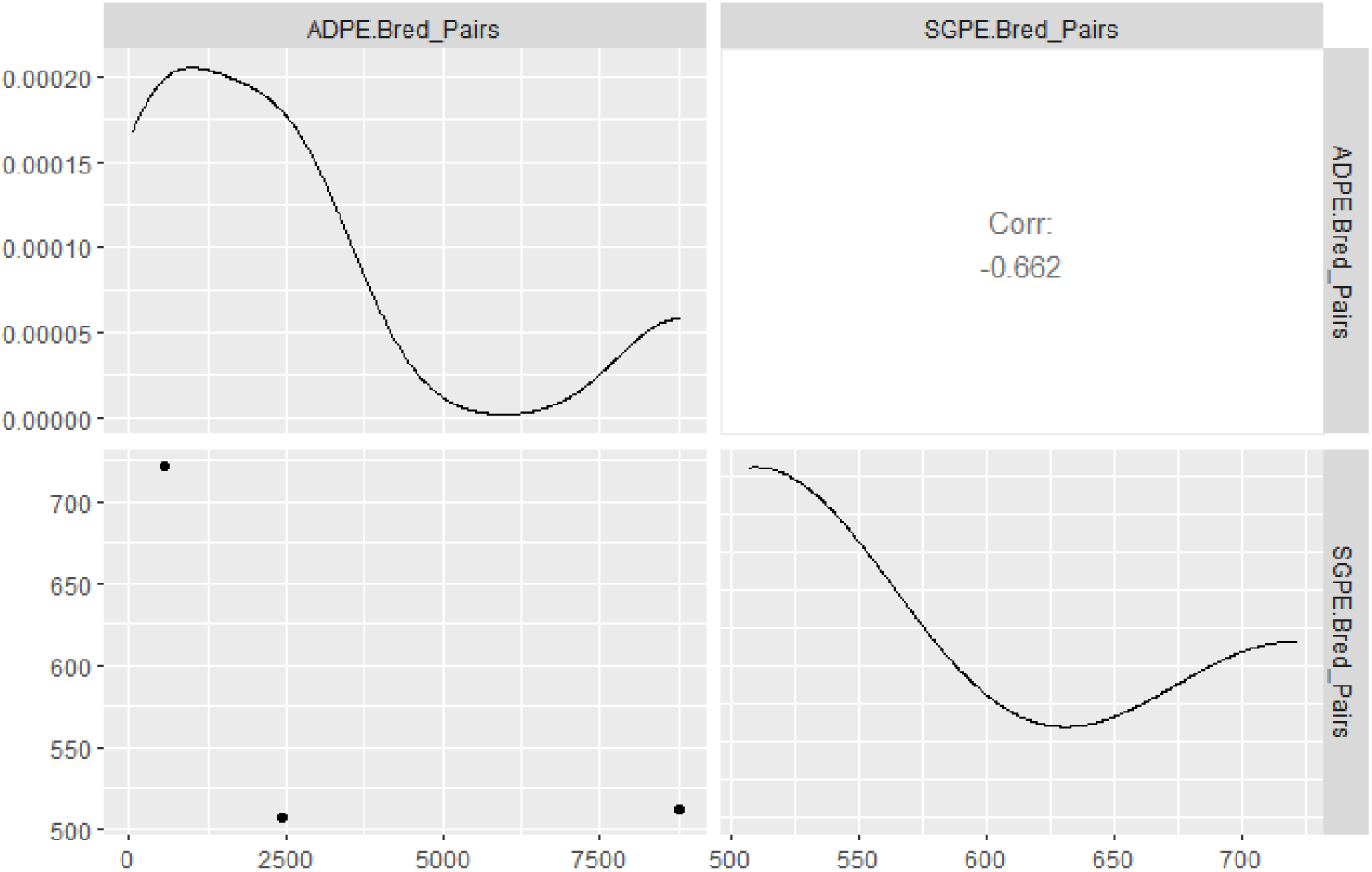
Pearson’s correlation coefficients (r) between Adélie (ADPE) and southern giant petrel (SGPE) breeding pairs from Penguin Island.

**Figure 2S:**
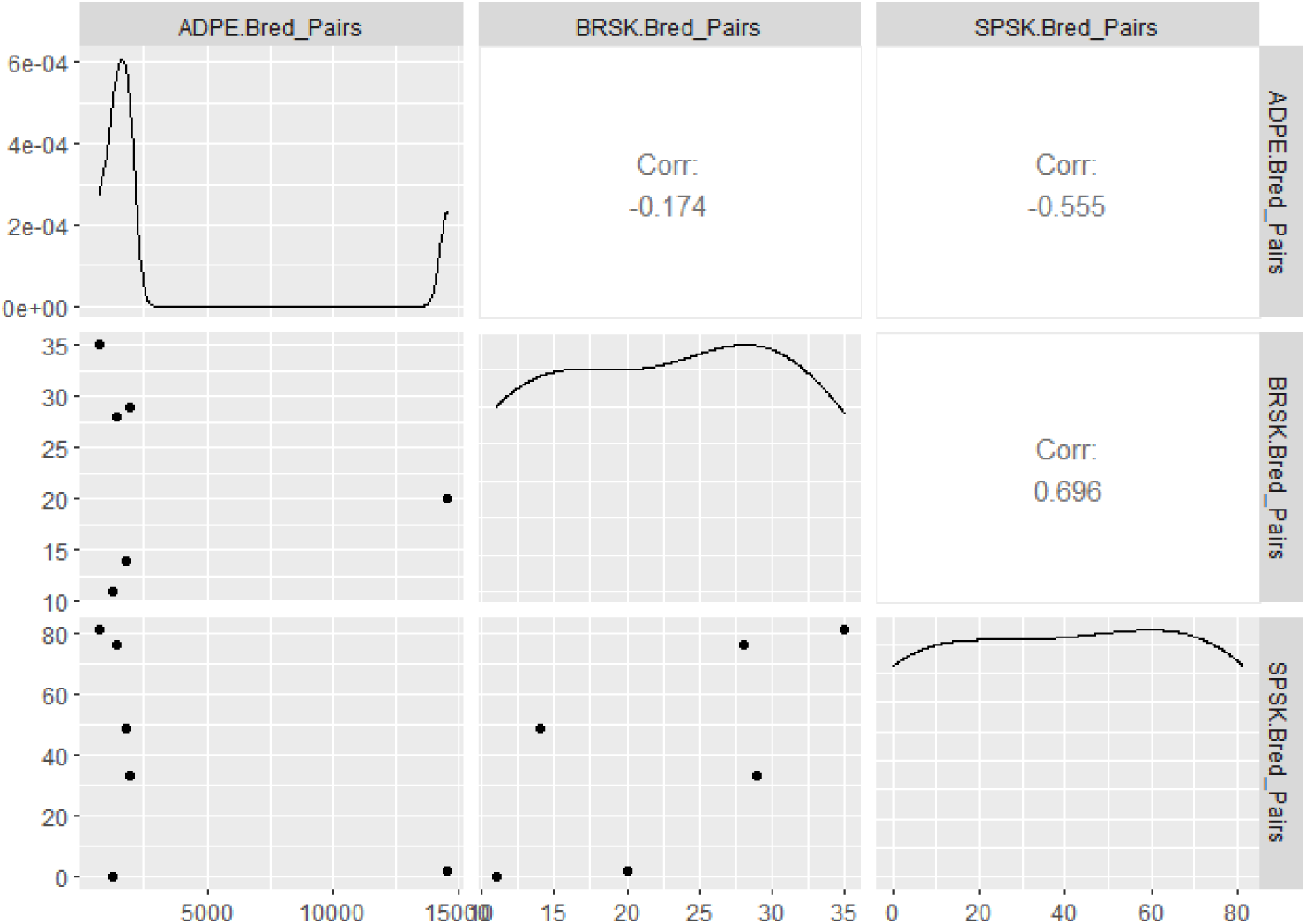
Pearson’s correlation coefficients (r) between Adélie (ADPE), brown skua (BRSK) and south polar skua (SPSK) breeding pairs from Potter’s Peninsula.

**Figure 3S:**
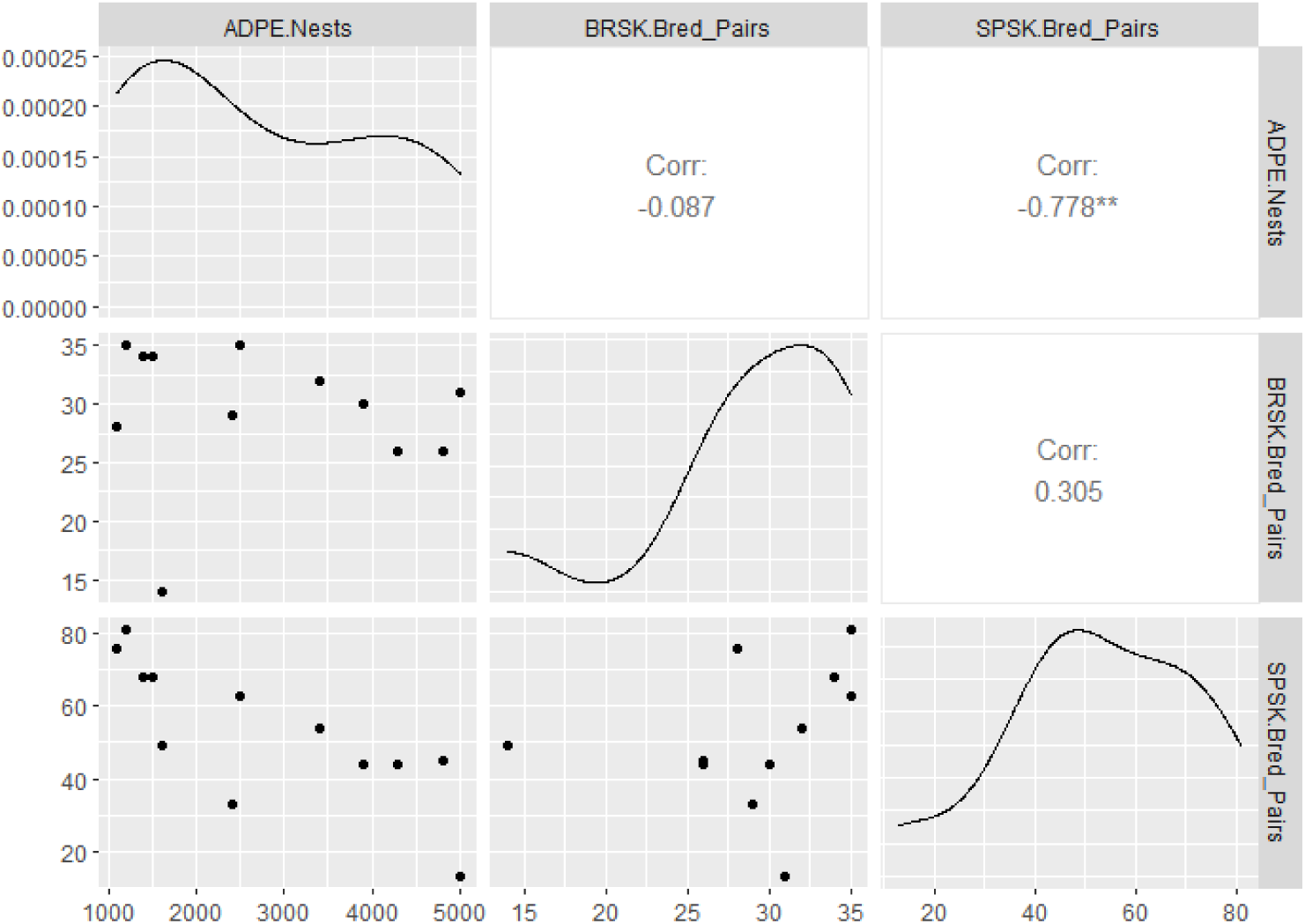
Pearson’s correlation coefficients (r) between Adélie (ADPE), brown skua (BRSK) and south polar skua (SPSK) breeding pairs from Potter’s Peninsula.

**Figure 4S:**
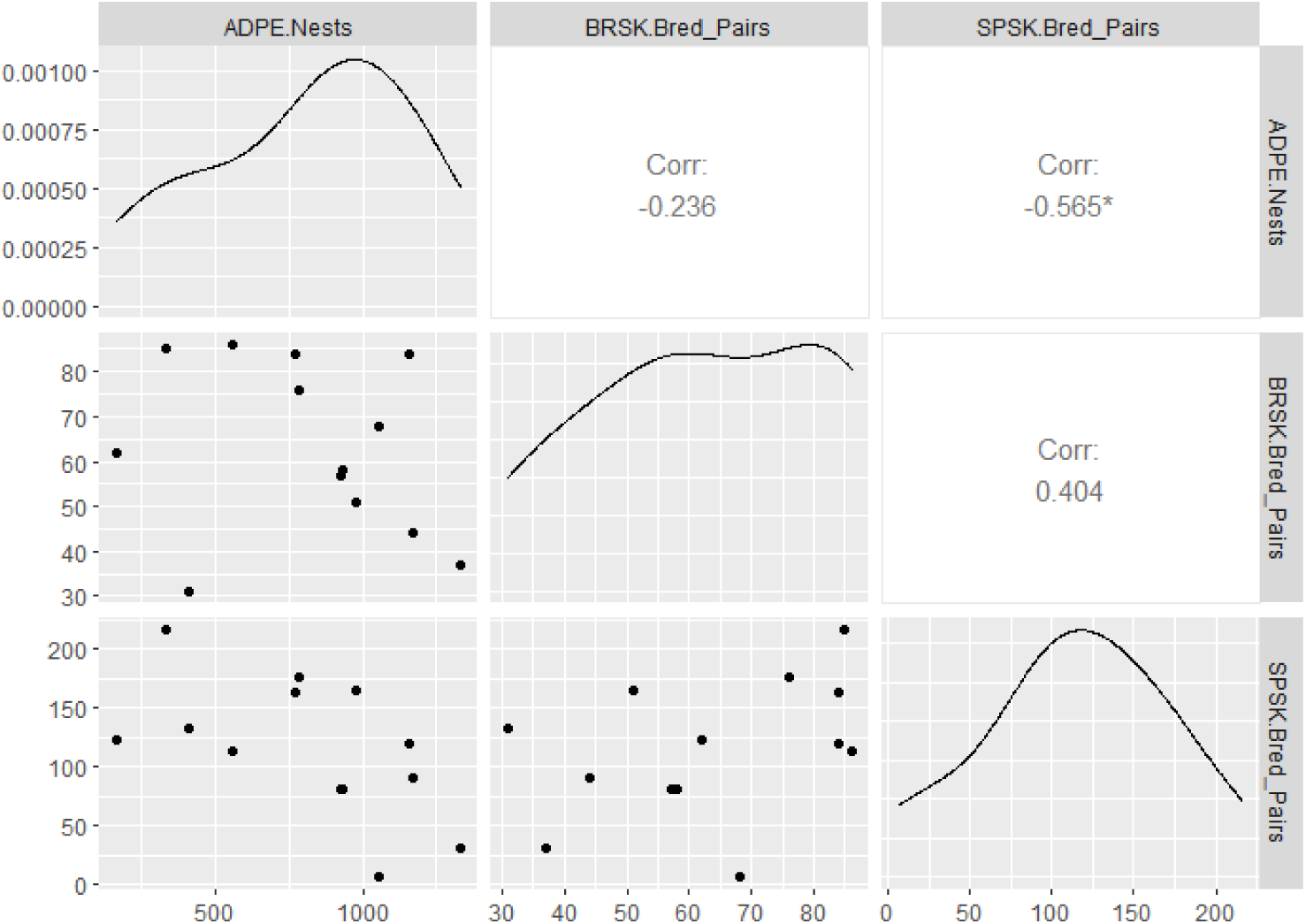
Pearson’s correlation coefficients (r) between Adélie (ADPE) nests and brown skua (BRSK) and south polar skua (SPSK) breeding pairs from Ardley Island.

**Figure 5S:**
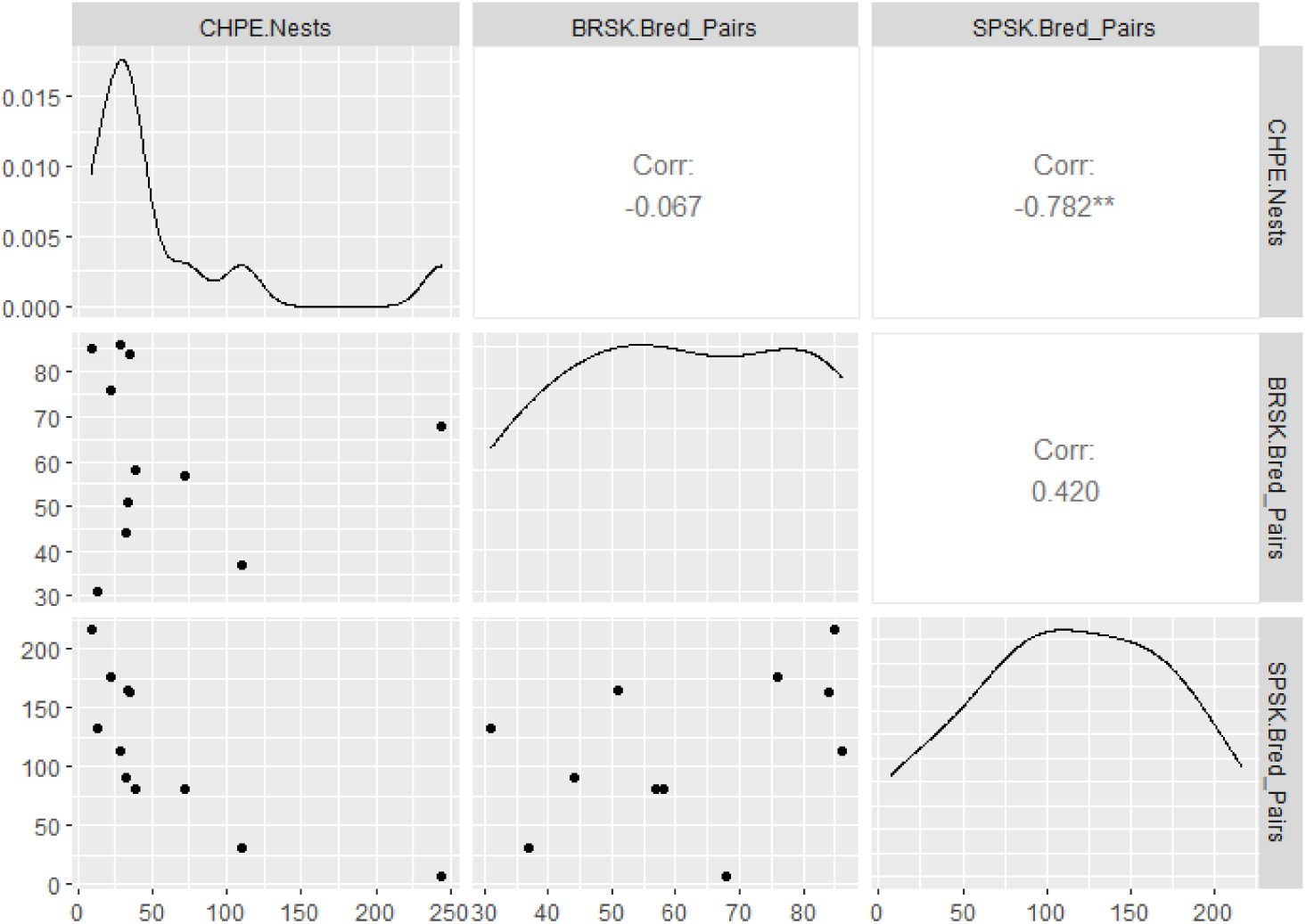
Pearson’s correlation coefficients (r) between chinstrap penguin (CHPE) nests, brown skua (BRSK) and south polar skua (SPSK) breeding pairs from Ardley Island.

**Figure 6S:**
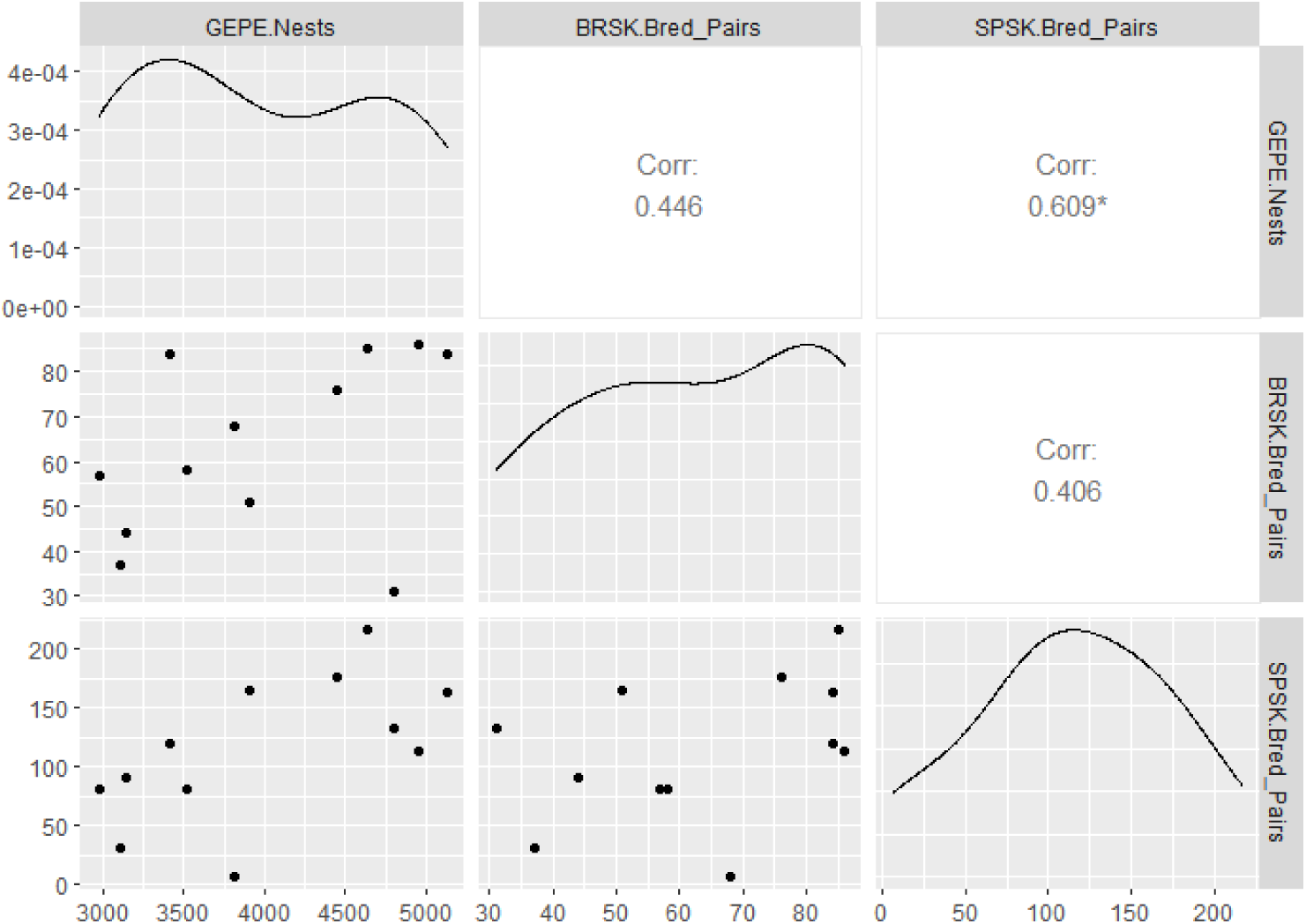
Pearson’s correlation coefficients (r) between gentoo penguin (GEPE) nests, brown skua (BRSK) and south polar skua (SPSK) breeding pairs from Ardley Island.

**Figure 7S:**
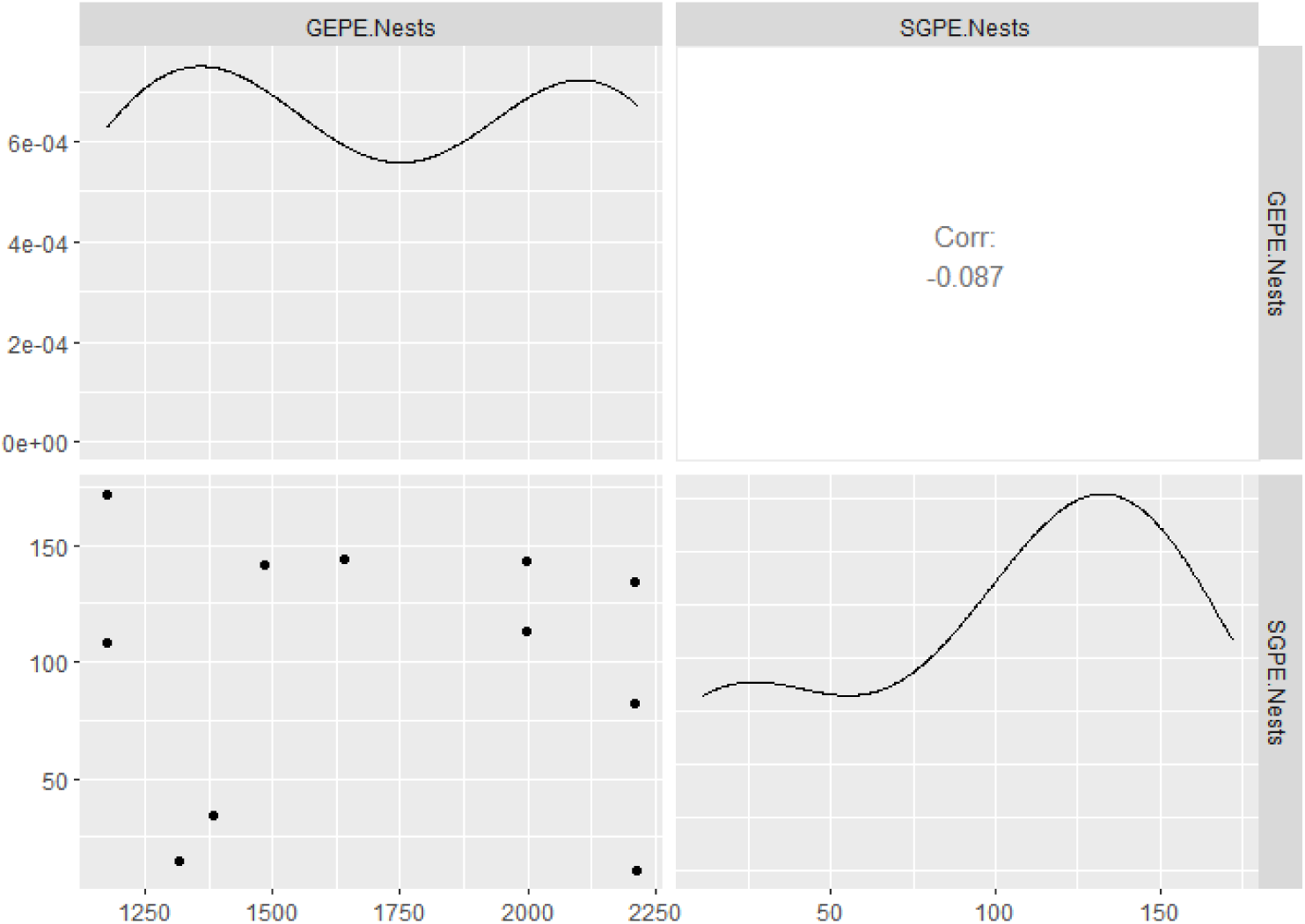
Pearson’s correlation coefficients (r) between gentoo penguin (GEPE) and southern giant petrel (SGPE) nests from Barrientos Island.

**Figure 8S:**
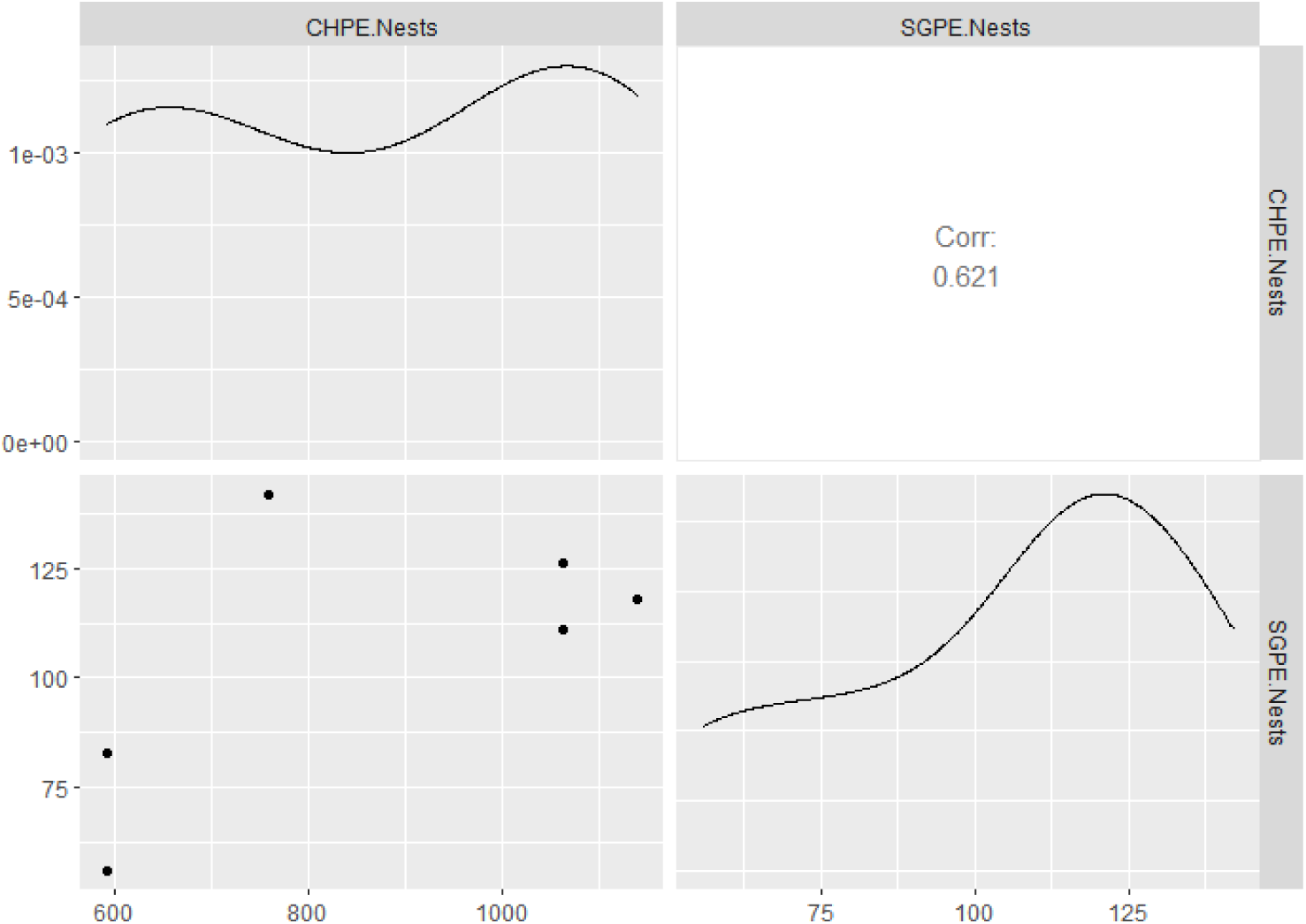
Pearson’s correlation coefficients (r) between chinstrap penguin (CHPE) and southern giant petrel (SGPE) nests from Hannah Point.

